# Metronomic therapy prevents emergence of drug resistance by maintaining the dynamic of intratumor heterogeneity

**DOI:** 10.1101/2021.01.04.425214

**Authors:** Maryna Bondarenko, Marion Le Grand, Yuval Shaked, Ziv Raviv, Guillemette Chapuisat, Cécile Carrère, Marie-Pierre Montero, Mailys Rossi, Eddy Pasquier, Manon Carré, Nicolas André

**Author notes:** Correspondence and requests for materials should be addressed to: Manon Carré, PhD, Cancer Research Center of Marseille, Faculty of Pharmacy, Bd Jean Moulin 13885 Marseille Cedex 5, France, Nicolas André, MD, PhD, Cancer Research Center of Marseille, Faculty of Pharmacy, Bd Jean Moulin 13885 Marseille Cedex 5, France. These authors contributed equally to this work (Maryna Bondarenko, Marion Le Grand). **Classification** Major classification: Biological sciences Minor classification: Computational and System biology.

## Abstract

Despite recent advances in deciphering cancer drug resistance mechanisms, relapse is a widely observed phenomenon in advanced cancers, mainly due to intratumor clonal heterogeneity. How tumor clones progress and impact each other remains elusive. By better understanding clone dynamics, we could reveal valuable biological insights and unveil vulnerabilities that could be therapeutically exploited. In this study, we developed 2D and 3D non-small cell lung cancer co-culture systems and defined a phenomenological mathematical model. Our results demonstrated a dominant role of the drug-sensitive clones over the drug-resistant ones under untreated conditions. Model predictions and their experimental *in vitro* and *in vivo* validations indicated that metronomic schedule leads to a better regulation of tumor cell heterogeneity over time than maximum-tolerated dose schedule, while achieving control of global tumor progression. We finally showed that drug-sensitive clones exert a suppressive effect on the proliferation of the drug-resistant ones through a paracrine mechanism way, which is linked to metabolic cell clone activity. Altogether, these computational and experimental approaches allow assessment of drug schedules controlling drug-sensitive and -resistant clone balance and highlight the potential of targeting cell metabolism to manage intratumor heterogeneity.

**Significance:** Combined computational and experimental models reveal how drug-sensitive tumor cells exert their dominance over drug-resistant cells and how it impacts optimal chemotherapy scheduling.

## Introduction

The advent of genomic medicine has increased our appreciation of intratumoral heterogeneity, which refers to clonal diversity among the tumor cells of a single patient. Even after malignant transformation, a cancer remains dynamic. This ongoing evolution generates a heterogeneous tumor mass consisting of cancer cells harboring distinct molecular signatures (1). Intratumor heterogeneity has emerged as a key mechanism that underlies therapeutic resistance and constitutes a genuine challenge for oncologists (2,3). Drug administration has, in itself, direct phenotypic consequences on tumor behavior and the emergence of resistant clones (4). Selective pressures exerted by anti-cancer drugs can lead to the expansion of resistant clones that either existed before the onset of therapy or that emerged as a result of treatment administration (2,3). Deciphering how tumor clones progress and impact each other could provide valuable biological insights and unveil vulnerabilities that could be therapeutically targeted.

Most research efforts are still focused on treatments that maximally kill tumor cells while minimizing toxicity to the host. Conventional chemotherapeutic drugs are currently predominantly administrated at or near the maximum tolerated dose (MTD), which gives the largest possible amount of drug at the beginning of the cycle and then let the patient recover from toxicities. However, it has been demonstrated that a cancer clone competition occurs within each individual tumor with the drug sensitive subpopulation of tumor cells proliferating at the expense of the resistant phenotype cells (5). In this context, the traditional MTD protocol leads to the elimination of chemosensitive clones, making way for the emergence of resistance clones (6,7). Modulating the dose and frequency of chemotherapy administrations in order to manage a constant tumor volume could bring substantial benefits over existing MTD protocols. Metronomic chemotherapy (MC) is defined as the chronic administration of chemotherapeutic agents at relatively low, minimally toxic doses, and with no prolonged drug-free breaks (8,9). Over the past two decades, MC protocols have shown promising results in several types of cancers, including ovarian, breast or lung cancers (10). A better understanding of the tumor dynamics under treatment is needed to design therapeutic strategies exploiting drug-resistant clones’ vulnerabilities.

In this study, we developed 2D and 3D heterogenous non-small cell lung cancer (NSCLC) co-culture models and phenomenological mathematical model to better understand intratumor heterogeneity. We revealed for the first time that metabolic activity of drug-sensitive clones could play a key role in controlling the proliferation of drug-resistant clones through a paracrine fashion. Moreover, metronomic schedule was predicted as the best strategy to control tumor volume without emergence of resistant clones.

## Materials and Methods

### Cell lines

The NSCLC cell lines used were A549 (RRID:CVCL_0023), A549/EpoB40 (RRID:CVCL_4Z15, epothilone B highly resistant cells derived from A549, kindly gave by Horwitz SB, AECOM, NY, USA), A549/VP16 (etoposide resistant cells derived from A549, kindly gave by Morjani H, Reims, France), H1975 (RRID:CVCL_1511), H1650 (RRID:CVCL_1483) and HCC827 (RRID:CVCL_2063) cell lines and the human colon adenocarcinoma cell lines were HT29 (RRID:CVCL_0320) and HT29/Rox1 (oxaliplatin resistant cells derived from HT29, kindly gave by Leloup L, INP, Marseille, France). The human NSCLC cell lines were grown in RPMI-1640 medium (Gibco, Thermo Fisher Scientific, France) and the human colon adenocarcinoma cell lines were cultured in DMEM medium (Gibco, Thermo Fisher Scientific, France). All the media were supplemented with 10% fetal bovine serum (FBS), 1% glutamine and 1% penicillin streptomycin (Gibco, Thermo Fisher Scientific, France). Patupilone 40 nM was added to RPMI-1640 medium for routine A549/EpoB40 cell culture and for cell adhesion. NSCLC cell lines were transfected by the mitochondrion-targeted DsRed (mtDsRed, Clontech, California, USA) plasmid or the enhanced green fluorescent protein (EGFP) plasmid (pEGFP-C1, Clontech/Takara Bio Europe). Transfections were achieved using Lipofectamine 2000 (Invitrogen, Thermo Fisher Scientific, France) according to the protocol supplied by the manufacturer. Stable cell lines were obtained after geneticin selection (800 μg/mL; Thermo Fisher Scientific, France). Cell sorting (FACScan, BD Biosciences) was also performed systematically to obtain high quality fluorescence cells. Cells were maintained in culture at 37°C with 5% CO_2_ and regularly screened to ensure the absence of mycoplasma contamination (MycoAlert, Lonza).

### Drugs and Reagents

Patupilone (epothilone B, Novartis, France), etoposide and cisplatin (Mylan, PA, USA), oxaliplatin (Sanofi-Aventis, France) and FX11 (cat. no. 427218; Merck Millipore, Lyon, France) were prepared in dimethylsulfoxide (DMSO, VWR), and were freshly diluted in the culture medium for experiments. The highest concentration of DMSO to which the cells were exposed was 0.001%.

### Establishment of homo- and heterogeneous 2D co-culture models

To yield experimental heterogeneous adherent cell co-cultures, 4.6×10^4^ drug-sensitive (A549-DsRed or HT29-DsRed) cells and 1.10^4^ drug-resistant (A549/EpoB40-GFP, A549/VP16-GFP or HT29/Rox1-GFP) cells were simultaneously seeded in a 6-well plate. Drug-sensitive A549 and drug-resistant A549/EpoB40 cells were first seeded in their specific culture medium. The two cell lines were initially separated by a silicon insert that was removed 24h after seeding. The homogeneous 2D co-cultures, which serve as control, included either A549-DsRed with A549-GFP or A549/EpoB40-DsRed with A549/EpoB40-GFP, in the same proportions as in the heterogeneous structures. The culture medium was changed 5 times a week in non-treated conditions and for cells exposed to MC regimens (0.5 nM patupilone). In the MTD conditions, cells were exposed to drug (5 nM patupilone) once a week for 24 hours, and incubated in daily-changed drug-free culture medium the four other days. To analyze the sensitive and resistant cell growth over time, DsRed and GFP fluorescent signals were recorded with a well-scanning mode microplate reader (PHERAStar FS, BMG LABTECH). Images were acquired by a Leica DM-IRBE microscope (Leica Microsystems). Cell proliferation was expressed as a percentage of the fluorescence signal measured at day 0.

### Establishment of homo- and heterogeneous 3D co-culture models

To obtain heterogeneous co-culture spheroids in 96-well round bottom plates, drug-resistant (A549/EpoB40-DsRed or HT29/Rox1-DsRed) cells were resuspended with drug-sensitive (A549-GFP or HT29-GFP) cells at the same ratio than the 2D co-culture models in a volume of 100μL culture media supplemented with 20% of methylcellulose (Sigma-Aldrich). The control 3D structures (homogeneous co-cultures) included A549/EpoB40-DsRed with A549/EpoB40-GFP cells, in the same proportions as the heterogeneous structures. After 48h, spheroids were treated following MC (0.5 nM patupilone or 0.2 μM oxaliplatin) and MTD treatment schedules (5 nM patupilone or 2 μM oxaliplatin). Resistant cell growth within the 3D micromasses was daily analyzed by recording the DsRed fluorescent signals as described for 2D co-cultures. Each signal was then converted into a number of resistant cells present in the structure, *via* a range of calibration carried out previously with 3D spheroids made of 500 to 150,000 A549/EpoB40-DsRed cells.

### Cell viability Assay

Cells were seeded in 96-well plates to be treated during 72 hours with MTAs. Cell survival was measured by using the colorimetric MTT assay (Sigma-Aldrich) as we previously performed (11).

### Proliferation assay

A549/EpoB40 cells were grown in 96-well plates for 24h and then, they were exposed daily to conditioned media from A549, H1975 or HCC825 cultures previously treated or not-treated with FX11. Conditioned media from A549/EpoB40 cells were used as control. Conditioned media were obtained from cells seeded in 6-well plates. Before treatment cell conditioned media were centrifuged 30min at 10,000 × g at 4°C. At different indicated time points, cells were fixed with 1 % glutaraldehyde and stained with a 1 % Crystal-violet solution in 20 % Methanol. The stain was eluted with DMSO and absorbance was measured at 600nm with a Multiskan Ascent plate reader.

### Impedance measurements

The label-free dynamic monitoring of cell proliferation and viability in real time was done by using the xCELLigence technology of the RTCA SP system (ACEA Biosciences, Ozyme, France). Cells were seeded in 96-wells microplates (E-Plates). Attachment of the cells was followed every 5min over 24 hours, and cell proliferation was then monitored daily every 15 min until cell confluency. Cell-sensor impedance was expressed as cell index values that were normalized using the RTCA Software 2.0 to the first measurement after starting treatment.

### Transwell co-culture system

A 24-Multiwell Insert System with 1 μm pore size were used (BD Falcon, BD Biosciences, France). Cells were seeded at 1,200 cells in both upper and lower chambers, to ensure a sustained exponential growth for 10 days. Cell culture media were changed in both upper and lower chambers every day, in agreement with 2D and 3D co-culture experiments. Two experimental seeding conditions were tested: 1) A549-DsRed in the upper chamber and A549/EpoB40-GFP in the lower chamber; 2) A549/EpoB40-DsRed in the upper chamber and A549-GFP in the lower chamber. A549/EpoB40-DsRed and A549/EpoB40-GFP in the upper and lower chambers respectively served as control. Fluorescence microscopy was used to ensure that no cell migrated from one compartment to the other. Similar experiments were also conducted with H1650, H1975 or HCC825 seeded in the upper chamber and with A549/EpoB40 cells seeded in the lower chamber. At day 3 and day 7, cell proliferation in the wells and the inserts were determined by using crystal violet staining as described in proliferation assay part.

### Extracellular vesicles isolation

Extracellular vesicles (EVs) were isolated from supernatant of A549 and A549/EpoB40 cells by differential centrifugations. Cell culture medium was centrifuged at 100,000 × g for 3h. Soluble fractions were filtered through a 0.22μm sterile filter and then mixed with serum free medium. Cells were seeded in T25 flask and incubated overnight. Cell conditioned media were replaced by EVs depleted cell culture medium at 24, 48 and 72h. Cell supernatants were collected at different time points and centrifuged 30min at 10,000 × g at 4°C to eliminate cell debris (centrifuge CT15RE, VWR), then at 100,000 × g for 3h to eliminate larger vesicles (L7-55 Ultracentrifuge, Beckman Coulter). The pellet was washed with PBS and ultracentrifuged at 100,000 × g 1h at 4°C to obtain small vesicles (TL-100 Ultracentrifuge, Beckman Coulter). The final pellet was resuspended in PBS for subsequent experiments. EVs were fixed with 2% uranyl acetate and were visualized by using transmission electron microscopy to quantify their sizes (JEOL 1220, Japan). To evaluate involvement of A549 total and fractionated conditioned media on A549/EpoB40 cell vialibility, A549 and A549/EpoB40 cells were seeded in 96-well plates (2,500 cells/well). At 24, 48 and 72h cells were treated with total cell-free supernatant, EVs and soluble fraction from A549 and A549/EpoB40 cells (the last ones used as control). Cell viability was measured using MTT assay at different time points as described in cell viability assay part.

### Real-time metabolic analysis

Multiparameter metabolic analysis of intact cells was performed in the Seahorse XF24 extracellular flux analyzer (Seahorse Bioscience, Billerica, MA, USA) as previously described (12). Briefly, A549 and A549/EpoB40 cells were seeded in Seahorse XF24 plates (1,500cells/well) and incubated overnight at 37°C in 5% CO_2_. To normalize OCR and ECAR data to cell number, A549 and A549/EpoB40 cells were simultaneously seeded in a second multi-well plate for 24h and then were stained with crystal violet as described in proliferation assay part.

### Glucose consumption and lactate production assay

YSI 2950 (Life Sciences, USA) was used to measure the total flux of glucose and lactate as previously described (12). At each time point, media were collected, centrifuged 5min at 1,200rpm and kept at −20 °C before measurement. Metabolite concentrations were normalized to cell number as described for real-time metabolic analysis.

### *In vivo* studies

Eight weeks old male NOD SCID mice, generously obtained from the laboratory of Prof. Israel Vlodavsky (Rappaport Faculty of Medicine, Technion), were injected subcutaneously to the right flank with A549-DsRed cells together with A549/EpoB40-GFP cells, in a ratio of 7:3 (sensitive: resistant) respectively. Tumors were allowed to grow and when reached the size of 200mm^3^, mice were randomly divided into three groups and treated with (i) MTD paclitaxel (25mg/kg, intraperitoneally, once every 3 weeks); (ii) MC paclitaxel (1.2mg/kg, intra-peritoneally, daily); or (iii) vehicle control. Tumor growth was monitored twice a week using a caliper. Mice were sacrificed at day 26 post-treatment initiation. Tumors were then removed, and single cell suspensions were produced as previously described (13). Vehicle-treated tumors were also sampled at the size of 100 and 200mm^3^. Cells were then incubated with APC-conjugated anti-human HLA antibody (clone W6-32; cat. no. 311410; BioLegend, San Diego, CA). Cells were subsequently analyzed by flow cytometry for GFP cells/DsRed cells (sensitive/resistant) composition of the tumor mass. Animal studies were performed in accordance with the Animal Care and Use Committee of the Technion-Israel Institute of Technology.

### Statistical analysis

Each experiment was performed at least in triplicate. Data are presented as mean ± S.E.M. Statistical significance was tested using unpaired Student’s t test. For experiments using multiple variables, statistical significance was assessed *via* two-way ANOVA. A significant difference between two conditions was recorded for \**p* < 0.05; \*\**p* < 0.01; \*\*\**p* < 0.001.

## Results

### Co-culture system demonstrates a balance between drug-sensitive and drug-resistant clones

Genomic diversity within single tumors has long been recognized. It is now well-known that drug-resistant clones co-exist with drug-sensitive clones in untreated tumors (14). In this study, we developed a simple co-culture method for adherent NSCLC cells to mimic intratumor heterogeneity *in vitro*, which allows long-term studies (up to 3 weeks). Using GFP and DsRed stably expressing cells, we were able to analyze over time the tumor subpopulations dynamic within these two-dimensional (2D) heterogeneous co-cultures by daily recording fluorescence signals (Supplementary Fig. S1). A549/EpoB40-GFP NSCLC cells, which are resistant to cisplatin and patupilone, and the parental drug-sensitive A549-DsRed NSCLC cells were used at the initial co-culture seeding ratio of 20/80 % respectively (Supplementary Fig. S2A). As shown in **Figs. 1A, B**, drug-sensitive A549 clones continuously proliferated in the whole well with an invasion of the drug-resistant cells’ initial seeding zone from day 6. In the meantime, the drug-resistant A549/EpoB40 clones did not expand and fluorescence analysis confirmed the decrease in the drug-resistant subpopulation, which was progressively substituted by the drug-sensitive clones (**Figs. 1A, B**). Similar results were obtained with other 2D co-culture models, including an etoposide resistant A549/VP16 NSCLC model and an oxaliplatin resistant HT29/Rox1 colorectal cancer model (Supplementary Figs. S2B, C). As drug-resistant cells are described as being less fit than the drug-sensitive cells due to the phenotypic cost of resistance (7), cell growth kinetic assays were performed in drug-sensitive A549 and drug-resistant A549/EpoB40 cells. Our results demonstrated that the doubling time difference between cell clones was less than 15% (Supplementary Fig. S2D). Moreover, to definitively ascertain the suppressive effect of the drug-sensitive clones over the proliferation of the drug-resistant ones, different cell clone ratios were used. As shown in **Fig. 1C**, in a homogenous co-culture, the A549/EpoB40 cell proliferation was constant over time until reaching well confluency. When drug-resistant A549/EpoB40 cells represented 40 or 80% of the initial co-culture seeding, a suppressive effect of the drug sensitive parental cell clones on the A549/EpoB40 cell proliferation was observed. Our data thus confirm that the inhibition of drug-resistant clone growth resulted from their co-culture interaction with the drug-sensitive clones. We then developed a NSCLC 3D spheroid co-culture model that better recapitulates cell-cell interactions in carcinomas. Fluorescence signals were converted to a number of drug-resistant cells using a calibration curve (Supplementary Fig. S2E). Consistent with the 2D co-culture results, fluorescence analysis demonstrated a stabilization of drug-resistant clone growth within hetero-spheroids, while A549/EpoB40 cells continuously proliferated within homo-spheroids (**Fig. 1D**). These results support the dominant role of the drug-sensitive A549 clones over the drug-resistant A549/EpoB40 clones in a 3D NSCLC co-culture system. Altogether, these results validate our 2D and 3D co-culture models, showing a dominant role of the drug-sensitive clones over the drug-resistant ones in a NSCLC model under untreated conditions.

**Figure 1.**
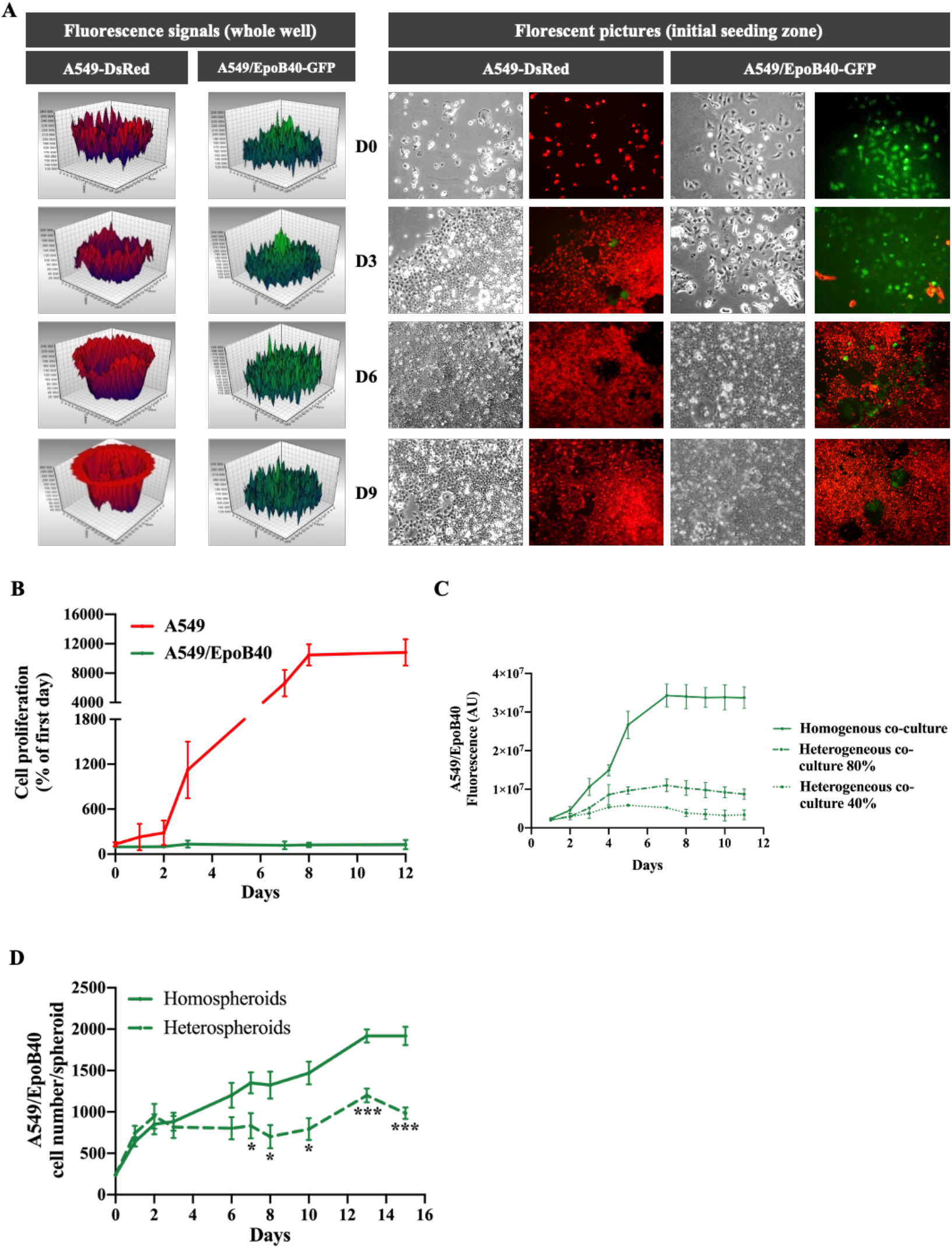
Drug-sensitive A549 clones inhibits the proliferation of the drug-resistant A549/EpoB40 clones. (A) Representative fluorescent signal using a well-scanning mode microplate reader (left panel) and microscopic pictures (right panel) of A549-DsRed and A549/EpoB40-GFP cells over time. (B) Cell proliferation of A549 and A549/EpoB40 by recording fluorescent signals (DsRed and GFP) over time. Data were expressed as a percentage of cell proliferation from day 0. (C) Fluorescent signal of A549/EpoB40 cells over time in homogenous or heterogenous 2D co-culture systems. (D) Relative A549/EpoB40 cell numbers in 3D hetero-spheroids and homo-spheroids measured by fluorescent signal over time. Data are shown as mean ± SEM. *, p < 0.05; ***, p < 0.001.

### Mathematical modeling predicts metronomic treatment as an optimal protocol to control intratumor heterogeneity

In order to better understand the influence of drug-sensitive clones over the drug-resistant ones, we have built a mathematical model, which is presented in details in Supplementary Fig. S3. It takes the following biological hypotheses into account:

1. In the well, there may be two types of cells denoted respectively drug-sensitive and -resistant cells.
2. In the absence of the other cell type, drug-sensitive and -resistant cells can grow freely with the same speed until the well becomes confluent.
3. All cells from both types are competing to colonize the available space in the well.
4. The presence of drug-sensitive cells in the well may have a suppressive role on the drug-resistant clone proliferation.

Our model was calibrated upon our experimental data carried out with A549 and A549/EpoB40 cells. Our first aim was to study whether a suppressive action of drug-sensitive clones over the resistant ones is actually required in order to explain our results presented above. First, we simulated that both cell clones were growing without a suppressive effect of the drug-sensitive cells (β = 0 in the model, Supplementary Fig. S3). As shown in **Fig. 2A**, our model predicts that both cell populations proliferate at the same speed until the well is confluent, maintaining the initial co-culture seeding ratio (20/80 respectively for drug-resistant and drug-sensitive clones). We then calibrated a suppressive effect of the drug-sensitive cells over the drug-resistant ones (β > 0 in the model). Our simulations suggest that the proliferation of drug-resistant cells decreases over time, while the drug-sensitive cells continuously grow until the well is confluent (**Fig. 2A**). These results demonstrated that our mathematical model fits better with our experimental data when a suppressive effect is taken into account, confirming the dominant role of the drug-sensitive clones over the drug-resistant ones in a NSCLC model.

**Figure 2.**
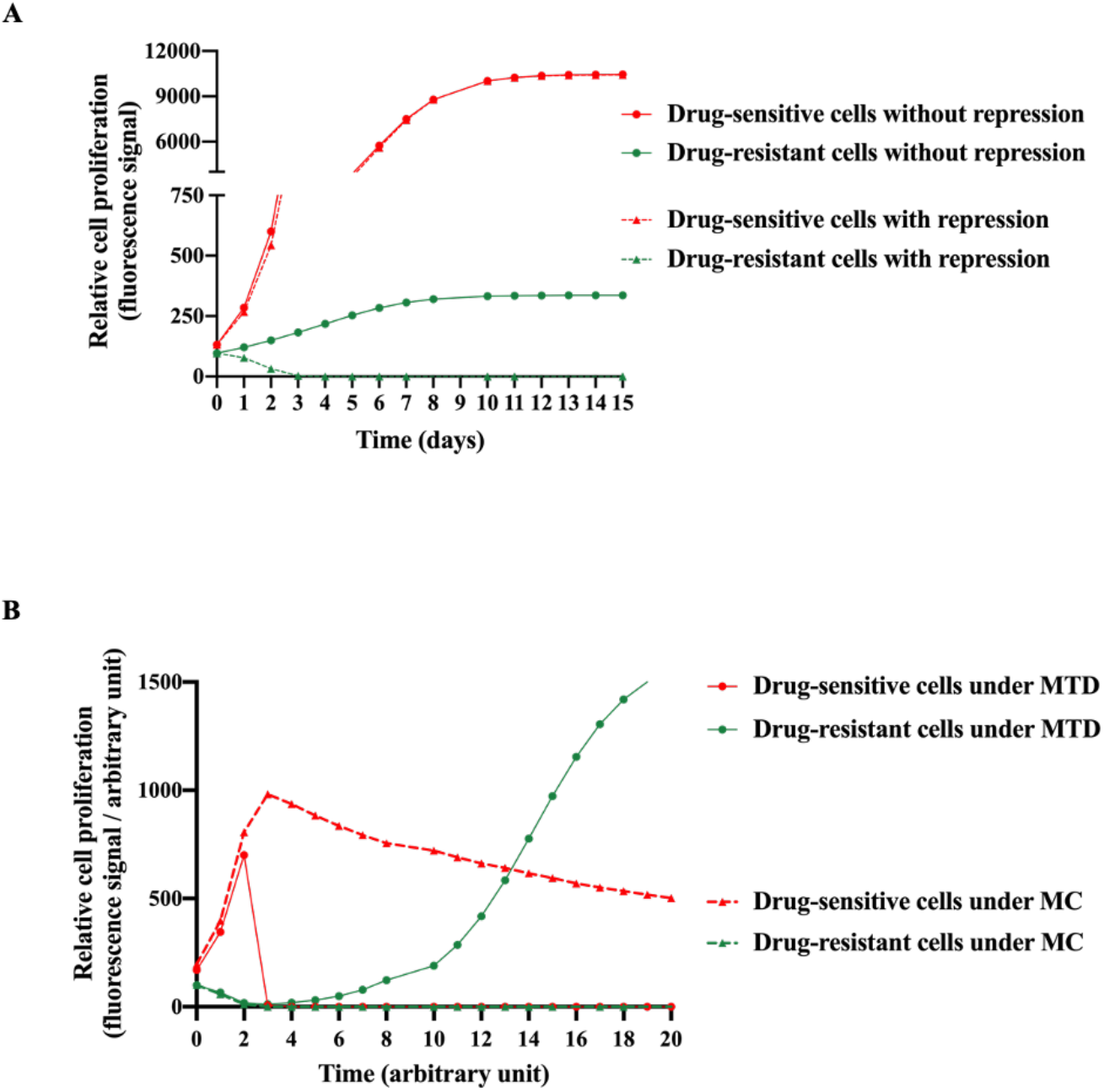
Model-based in silico simulation of the interaction between sensitive and resistant clones with or without treatment. (A) Model-based *in silico* simulations of relative drug-sensitive and drug-resistant cell proliferation over time with or without a repressive effect exerted by the drug-sensitive cells over the drug-resistant ones. (B) *In silico* simulations of relative drug-sensitive and drug-resistant cell proliferation under MC and MTD schedules and taking into account the suppressive effect of the drug-sensitive cells over the drug-resistant ones (β > 0 in the model).

In order to understand the influence of the suppressive action of sensitive cells upon the efficiency of a treatment, we then introduced the effect of chemotherapy in our mathematical model (Supplementary Fig. S3). The biological hypotheses considered are the following:

1. The chemotherapeutic agent only acts on drug-sensitive cells.
2. A delay between the injection of the chemotherapeutic agent and its impact on cell functions is taken into account.
3. The chemotherapeutic agent concentration is assumed constant in absence of an experimenter intervention.

Taking into account the suppressive effect of drug-sensitive clones over the drug-resistant ones, simulations of different dosing regimens were performed. Our model predicts that following a MTD treatment, the drug-sensitive cell proliferation is strongly inhibited, while the drug-resistant cell proliferation resumes after a certain delay until the well is confluent (**Figure 2B**). We then changed the schedule of administration using a metronomic protocol (MC). A long-term control of drug-sensitive cell proliferation was obtained, with no drug-resistant cell proliferation (**Fig. 2B**). The analysis of the mathematical model therefore suggests that a metronomic schedule should be more efficient than MTD-based conventional chemotherapy to maintain a heterogeneous tumor as small as possible.

### *In vitro* and *in vivo* biological validations of mathematical model predictions

On the basis of our *in silico* model predictions, we evaluated the antitumor efficacy of two treatment regimens in the 2D heterogenous co-culture system: a MTD-like treatment schedule based on the IC_50_ values of the drug-sensitive cells (5nM of patupilone, once a week for 24h) and a MC-like treatment schedule based on protracted low-dose drug administration (0.5nM of patupilone, five times a week). Daily fluorescence recording revealed that both treatment schedules led to a global decrease of tumor proliferation rate in two different ways. Indeed, MTD-like treatment schedule was highly effective in decreasing proliferation of the drug-sensitive A549-DsRed cells (**Fig. 3A**), while the MC-like treatment schedule only resulted in a limiting drug-sensitive A549-DsRed proliferation (**Fig. 3B**). Meanwhile, as predicted by our mathematical model, the drug-resistant A549/EpoB40-GFP clones started to grow from day 5 under the MTD-like treatment schedule, and proliferated until the surface of the well was fully confluent (**Fig. 3A**). In sharp contrast, the MC-like treatment schedule was able to persistently suppress the growth of the drug-resistance A549/EpoB40-GFP clones (**Fig. 3B**). Collectively, these data suggest that the metronomic schedule would better manage drug-resistant clone proliferation than the MTD-like treatment schedule and validate our mathematical models. We were also able to predict a specific threshold of the chemotherapeutic agent concentration, called C_*threshold*_, which keep a small proportion of drug-sensitive cells for maintaining a repressive effect on the drug-resistant cell proliferation. Using our experimental data, our model has been calibrated and our results indicated that the C_*threshold*_ was at 0.6 nM (**Fig. 3C**).

**Figure 3.**
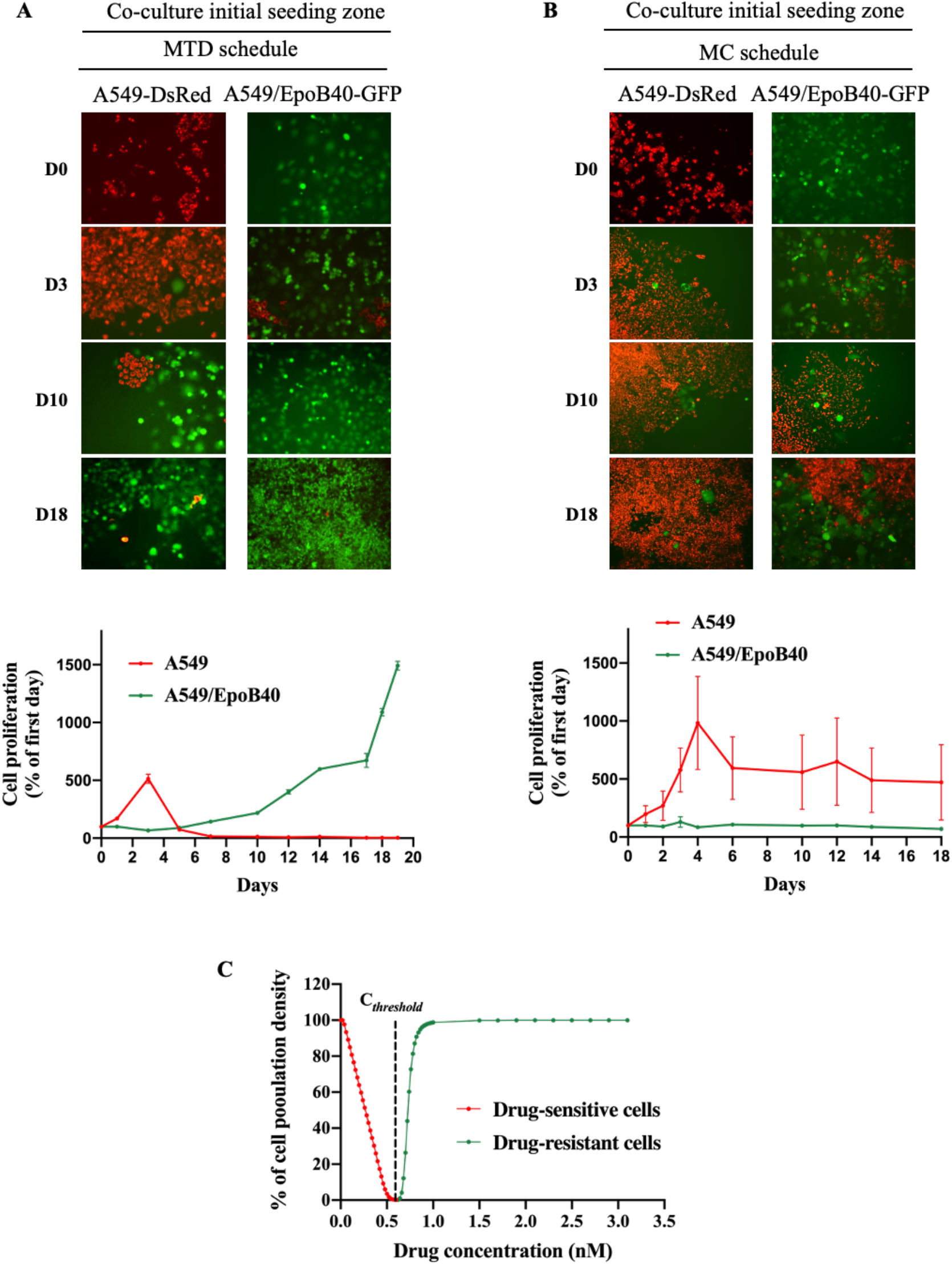
Metronomic schedule prevents from selecting drug-resistant clones in 2D NSCLC co-culture models. (A, B) Representative fluorescent pictures of A549-DsRed and A549/EpoB40-GFP cells over time under MTD (A) or MC (B) schedules. Cell proliferation of A549 and A549/EpoB40 by recording fluorescent signals (DsRed and GFP) over time under MTD (A) or MC (B) schedules. Data were expressed as a percentage of cell proliferation from day 0. (C) *In silico* simulations of the C_*threshold*_ allowing to control the proliferation of a heterogenous tumor.

We then confirmed our 2D heterogenous co-culture data and mathematical simulations in a 3D co-culture model. Our results demonstrated that the proliferation of the drug-resistant clones was partially repressed under the MC-like treatment in comparison to the MTD-like treatment (**Fig. 4A**). Indeed, while the drug-resistant cell proliferation increased by a factor 2.1 under the MC-like treatment, a fold-3 increase was observed under MTD-like treatment (**Fig. 4A**). Moreover, the MC-like treatment effect on repressing the drug-resistant population was also validated in the 3D HT29/Rox1 colorectal cancer model (**Fig. 4B**). While no difference was observed between the non-treated condition and metronomic schedule, the drug-resistant cell proliferation increased by a factor 2.7 under the MTD schedule (**Fig. 4B**). To validate our results in an *in vivo* model, a mixture of drug-sensitive A549-mtDsRed and drug-resistant A549/EpoB40-GFP cells (ratio of 7/3 – sensitive / resistant) was inoculated in the right flank of NOD SCID mice. To first ensure the constant cell heterogeneity of our model over time in the absence of treatment, we analyzed *ex vivo* the tumor cell composition at three different time points: when tumor size reached 100 and 200mm^3^ and, at study end point. As shown in **Fig. 4C**, the tumor cell composition did not change over time, confirming the effect of drug-sensitive clones in controlling the growth of their drug-resistant counterparts in untreated condition. An MTD-like (paclitaxel at 25 mg/kg, once every 3 weeks) or MC-like (paclitaxel 1.2 mg/kg, daily) treatment schedules were then administered to mosaic tumor-bearing mice. Both treatments had a significant repressive effect on tumor growth (−67 +/− 8 and −48 +/− 4 %, respectively with MTD- and MC-like treatment schedules in comparison to control; **Fig. 4D**). Although there was no significant difference between the MTD- and the MC-like treatments on the tumor growth at study end point, the ratio of drug-sensitive to drug-resistant cells in the tumor mass was significantly higher in the MC-like treated mice. Indeed, following the MC-like treatment, the ratio increased by 124 +/− 85 % in comparison to control, while the MTD-like treatment led to a decrease by 53 +/− 15 % in comparison to control (**Fig. 4E**). Collectively, these results validate our mathematical model predictions and indicate that metronomic schedule leads to a better control of NSCLC cell heterogeneity than MTD schedule over time, while achieving control of global tumor volume.

**Figure 4.**
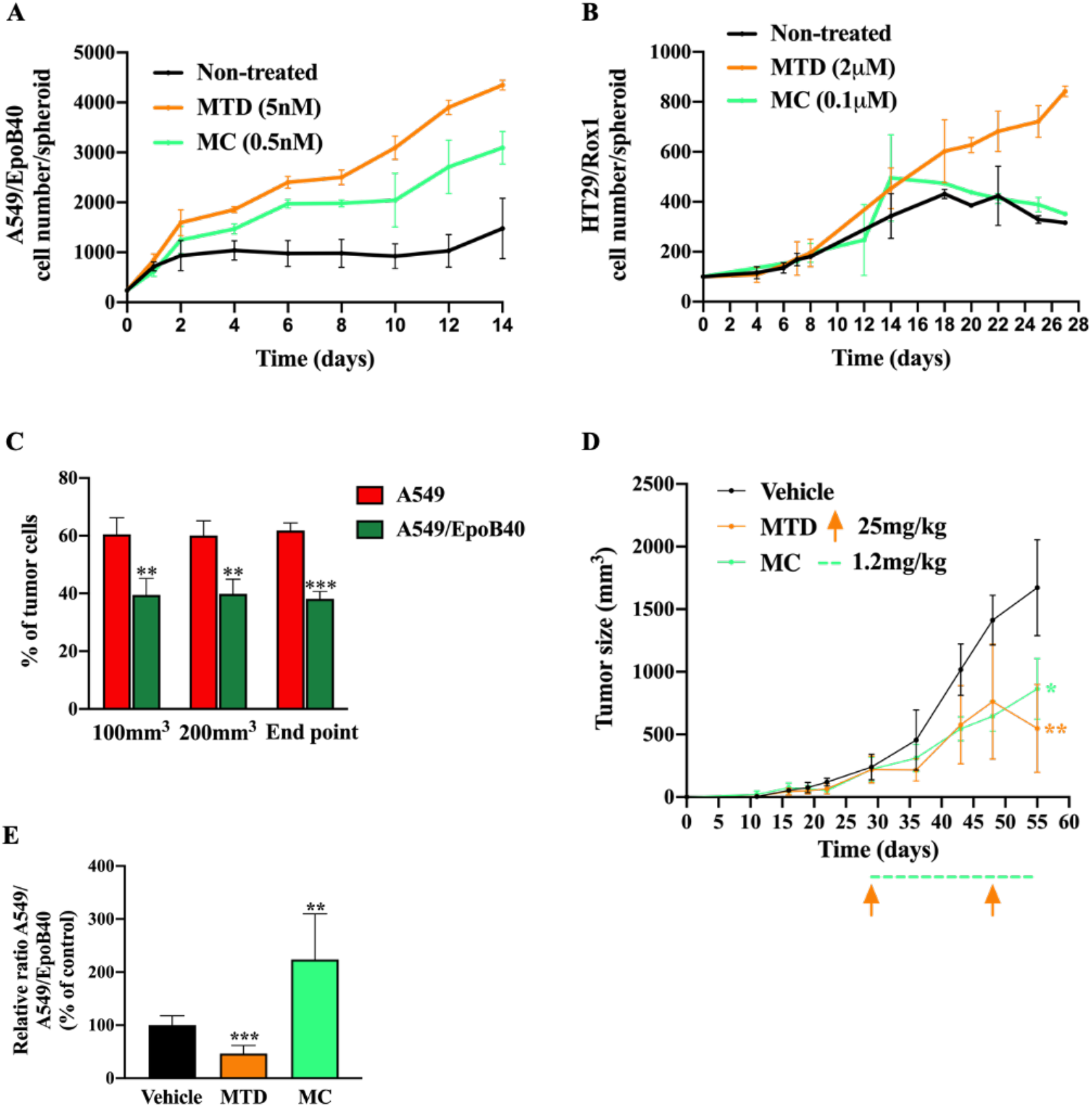
Biological validation of the mathematical model predictions in 3D spheroid and *in vivo* models. (A, B) Relative A549/EpoB40 (A) and HT29/rox1 (B) cell numbers in 3D hetero-spheroids measured by fluorescent signal over time under MTD (patupilone 5nM (A) or oxaliplatin 2μM (B)) or MC (patupilone 0.5nM (A) or oxaliplatin 0.1μM (B)) schedules. (C) *Ex vivo* tumor cell composition at three different time points in the untreated condition. Fluorescent cell signals were quantified by flow cytometry. (D) Tumor sizes were measured over time in untreated condition as well as under MTD (paclitaxel at 25 mg/kg, once every 3 weeks) and MC treatment (paclitaxel 1.2 mg/kg, daily). Significant differences compared to vehicle. Orange arrows represent MTD-drug administration, while dotted green line indicates the MC-drug administration period. (E) Relative ratio of the *ex vivo* tumor cell composition at the end point in untreated condition and following a MTD-like and MC-like treatment schedules. Fluorescent cell signals were quantified by flow cytometry. Data are shown as mean ± SEM. **, p < 0.01; ***, p < 0.001.

### Drug-sensitive clones control the proliferation of drug-resistant clones through a paracrine pathway

Many studies have demonstrated the role of paracrine pathways for cell-cell communication in cancer (15). To investigate whether the drug-sensitive NSCLC clones can exert their suppressive effect on the drug-resistant ones without direct cell interaction, we used Transwell^®^ systems of heterogeneous co-cultures. Our results demonstrated that in homogenous co-culture, A549/EpoB40 cells continuously grow over time (**Fig. 5A**). However, drug-resistant cell proliferation could be inhibited by indirect interaction with the drug-sensitive cells (**Fig. 5A**). This repressive effect was not specific to A549 cells as similar results were obtained with H1650, H1975 and HCC827 NSCLC cell lines (**Fig. 5A**). We further analyzed the effect of conditioned media using cell-free supernatants on the drug-resistant A549/EpoB40 cell growth by real-time impedance measurement. Our results showed a decrease in the proliferation of drug-resistant A549/EpoB40 cells treated with drug-sensitive A549 supernatants in comparison to treatment with the drug-resistant A549/EpoB40 supernatants (**Fig. 5B**, grey-shaded area). This was reflected by a significant drop in the A549/EpoB40 cell growth curve slope treated with A549 supernatant condition of 56 +/− 3 %, demonstrating a paracrine interaction between the two cell clones (**Figs. 5B, C**). Moreover, the discontinuation of exposure to drug-sensitive A549 supernatants led to a recovery of the drug-resistant A549/epoB40 cell proliferation (**Fig. 5B**, white-shaded area) as indicating with the same slopes for both cell populations (**Fig. 5C**). As it has recently become apparent that secreted extracellular vesicles (EVs) are proficient intercellular communication mediators (16), we separated vesicles from soluble fraction of drug-sensitive A549 cell-free supernatants by sequential centrifugations and analyzed their effect on drug-resistant A549/EpoB40 cell growth. The vesicles we extracted were exosomes based on their diameter, ranging from 50 to 200 nm as previously described (17) (data not shown). Our results showed that treatment with soluble fraction from drug-sensitive A549 cells significantly decreased the drug-resistant A549/EpoB40 cell viability over time as did A549 cell-free supernatant treatment in agreement with the impedance results (**Fig. 5D**). In comparison, EVs fraction did not significantly impact the drug-resistant A549/EpoB40 cell viability (**Fig. 5D**). Collectively, our data indicate that drug-sensitive A549 clones can control the proliferation of drug-resistant A549/EpoB40 clones through a paracrine pathway, independently of exosome secretion.

**Figure 5.**
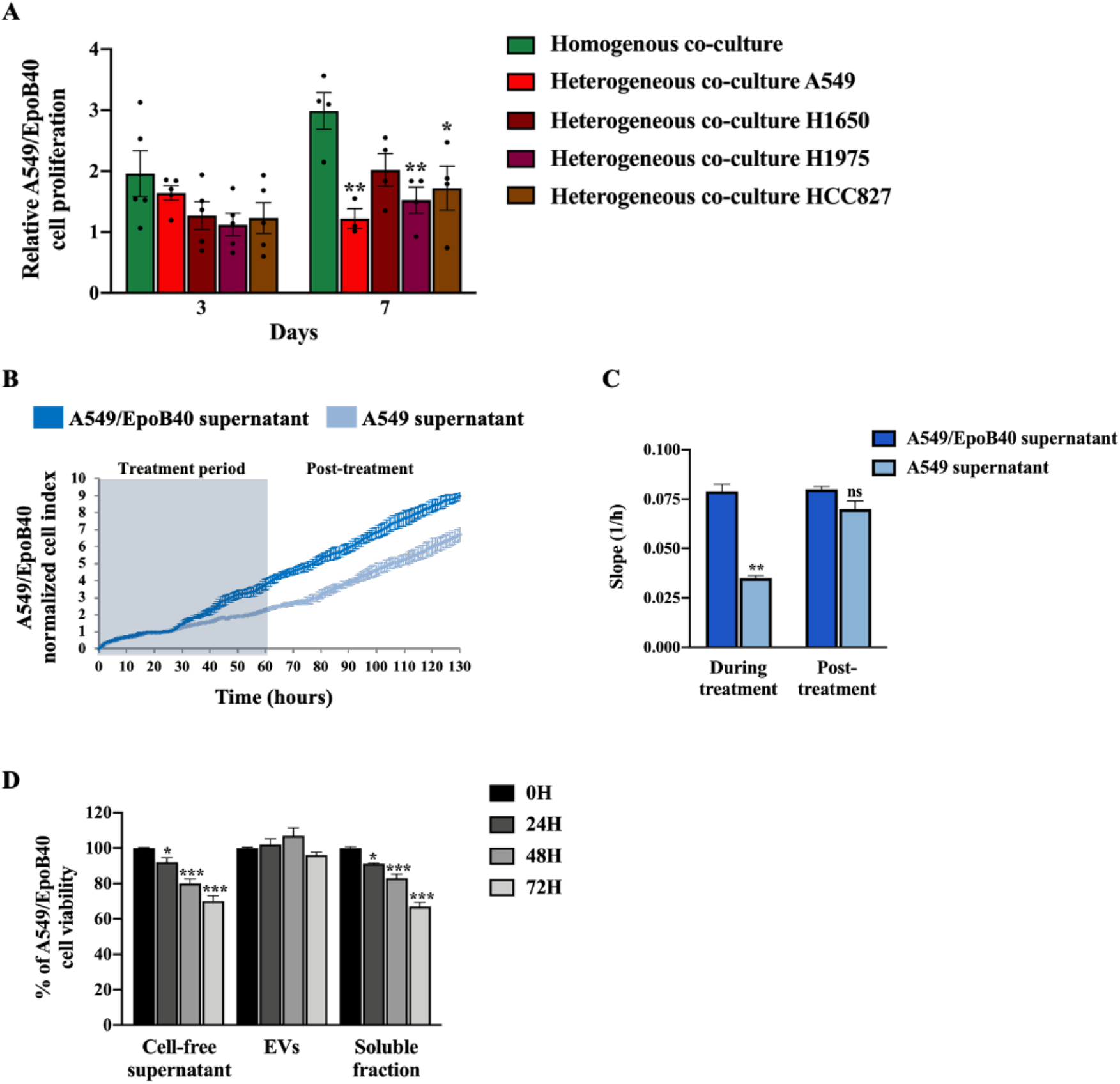
Drug-sensitive A549 clones control the proliferation of drug-resistant A549/EpoB40 clones through a paracrine pathway, independently of exosome secretion. (A) Relative quantification of A549/EpoB40 cell proliferation in homogeneous (A549/EpoB40 only) and heterogeneous (A549/EpoB40 with either A549, H1650, H1975 or HCC827) co-culture Transwell^®^. Cells were stained with crystal violet at indicated times. Data were expressed as a ratio of cell proliferation from day 0. (B) A549/EpoB40 cell growth was studied by real time impedance-based method. Twenty-four hours after seeding, A549/EpoB40 cells were treated daily with cell-free supernatants from A549 cultures and from A549/EpoB40 cultures that were used as control. Measurements were performed every 15 minutes. Cell index values were normalized to 24 h corresponding to the first measurement after starting treatment. Grey-shaded area indicates the duration of treatment. (C) Slopes were determined during the period of 48h supernatant treatment and 48h post-supernatant treatment. (D) Cell proliferatio of A549/EpoB40 cells measured by crystal violet assay after a 0, 24, 48 and 72h exposition to total cell-free supernatant, extracellular vesicles (EVs) or soluble fraction from A549 cultures. Results were expressed as a percentage of viability at T0. Data are shown as mean ± SEM. *, p < 0.05; **, p < 0.01; ***, p < 0.001.

### The metabolic activity of drug-sensitive clones is a key factor to maintain their repression on minority drug-resistant clones

It is now established that metabolic reprogramming happens in tumor tissue resulting in cancer cell phenotypes with different metabolic activities leading to adaptive/acquired resistance to anti-tumor therapy (18). To investigate whether the metabolic status of drug-sensitive A549 and drug-resistant A549/EpoB40 clones could be different, we performed real-time measurements of oxygen consumption rate (OCR) and extracellular acidification rate (ECAR), reflecting the mitochondrial and glycolytic activities, respectively. As illustrated in **Fig. 6A**, drug-resistant A549/EpoB40 cells had a higher overall mitochondrial metabolic efficiency than drug-sensitive A549 cells. Indeed, basal OCR increased by 85 +/− 7 % (p<0.001) in comparison to the drug-sensitive A549 cells. Moreover, maximal mitochondrial respiration, induced by FCCP, increased by 101 +/− 16 % (p<0.001) in comparison to the drug-sensitive A549 cells. Meanwhile, our results showed that drug-resistant A549/EpoB40 cells were not able to increase their glycolytic capacities following oligomycin administration, which suppresses mitochondrial energy production (**Fig. 6B**). This demonstrates that drug-resistant cells are more dependent on mitochondrial respiration than drug-sensitive cells, which mostly rely on glycolytic activities. As this type of altered energetic metabolism can result in the secretion of specific metabolites that can have a significant impact on tumor progression, dissemination and drug response (18), we next measured extracellular glucose and lactate concentration in cell-free supernatants. Our results showed that drug-sensitive A549 cells exhibited higher glucose consumption and lactate production than drug-resistant A549/EpoB40 cells. Indeed, at 48h, relative glucose and lactate concentrations were respectively 83 +/− 4 % lower and 93 +/− 15 % higher in drug-sensitive A549 cell-free supernatant in comparison to drug-resistant A549/EpoB40 cell-free supernatant (**Figs. 6C, D**). As our results suggest an important role of the glycolytic pathway in the drug-sensitive A549 cells, we focused on the potential role of LDHA (Lactate dehydrogenase A) being the primary metabolic enzyme that converts pyruvate to lactate in cancer cells (19) and a hallmark of aggressive cancers (20,21). We used FX11, a pharmacological inhibitor of LDHA at a non-cytotoxic concentration of 3μM (Supplementary Fig. S4A). Our results first showed that pre-treatment of drug-sensitive cells with FX11 was able to significantly reverse the suppressive effect of cell-free supernatants from A549, H1975 or HCC825 cells on the drug-resistant A549/EpoB40 cell proliferation (**Fig. 6E**, p<0.01). Our data also showed that supernatants from FX11 pre-treated drug-sensitive A549 cells restored the proliferative rate of drug-resistant A549/EpoB40 cells (**Fig. 6F**, Supplementary Fig. S4B). Altogether, our results suggest that the glycolytic activity of drug-sensitive A549 clones could play a major role on the drug-resistant A549/EpoB40 clone proliferation.

**Figure 6.**
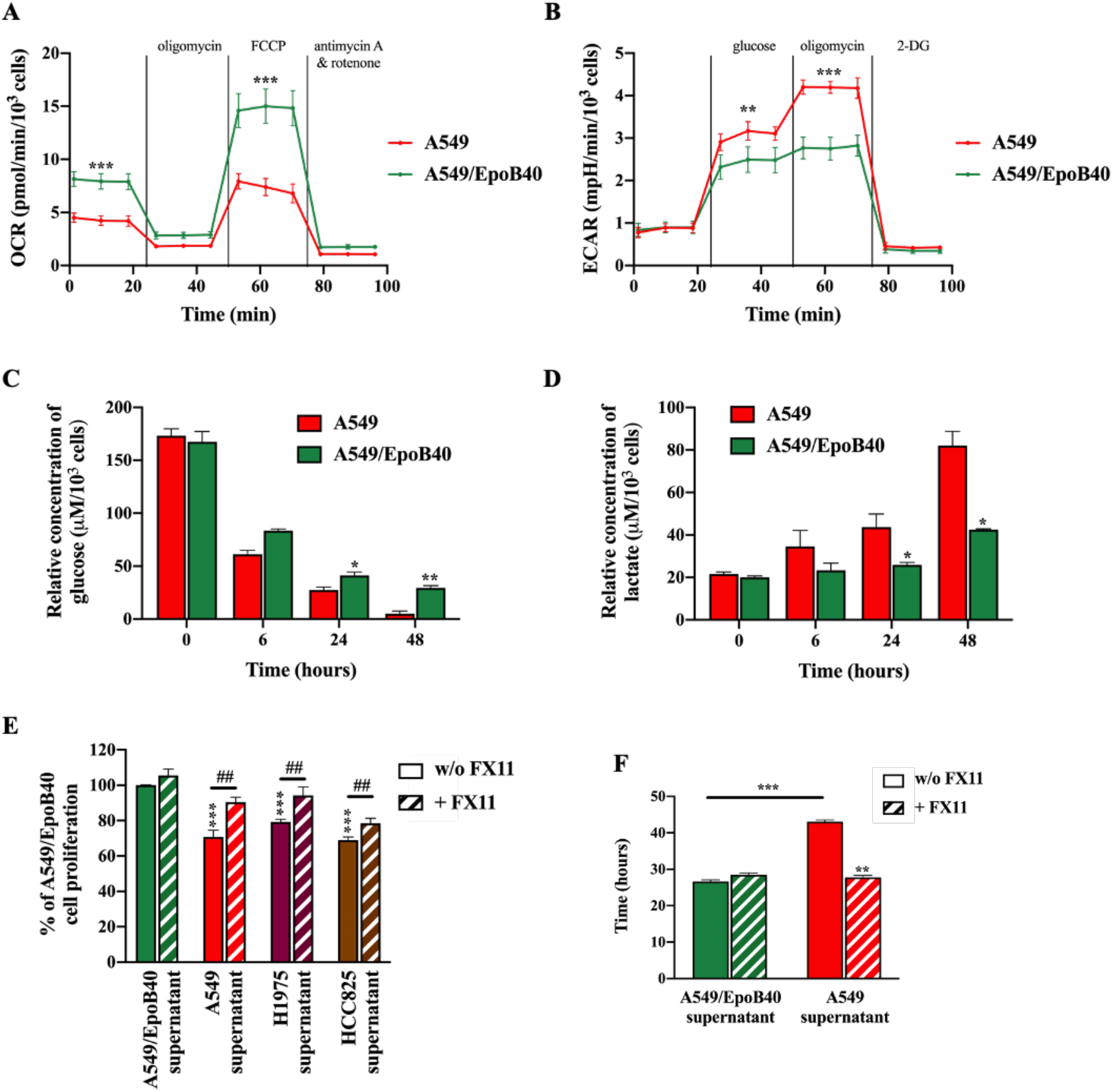
Drug-sensitive A549 clones have a higher glycolytic profile than drug-resistant A549/EpoB40 clones. (A, B) A549 and A549/EpoB40 cells were analyzed for (A) mitochondrial bioenergetics and (B) glycolysis using the Seahorse XF technology. (C, D) Supernatants from A549 and A549/EpoB40 were collected over time and analyzed for (C) glucose consumption and (D) lactate production by using the YSI 2900 instrument. (E) A549/EpoB40 cell proliferation was measured by crystal violet assay at 72 h after daily exposition to cell-free supernatants from A549, H1975 or HCC825 cultures previously incubated with or without 3μM of FX11. Supernatants from A549/EpoB40 cultures that were subjected to the same treatment were used as control. Results were expressed as a percentage of control cell proliferation. (F) A549/EpoB40 cell growth was followed by real time impedance-based method. A549/EpoB40 cells were exposed daily to cell-free supernatants from A549 and A549/EpoB40 cultures that were previously incubated with or without 3μM of FX11. Doubling time of A549/EpoB40 cells was calculated in different conditions. Data are shown as mean ± SEM. *, p < 0.05; **, p < 0.01; ***, p < 0.001.

## Discussion

Lung cancer is the leading cause of cancer-related death worldwide with non-small-cell lung cancer (NSCLC) being the most common type (22). Despite the emergence of targeted therapies and immunotherapies, chemotherapy is still the standard-of-care for NSCLC (23). Studies evaluating intratumor heterogeneity and cancer genome evolution in lung cancer patient cohorts have revealed that intratumor heterogeneity is a key factor contributing to the lethal outcome of lung cancer, therapeutic failure, and drug resistance (24,25). Here, we developed heterogenous NSCLC co-culture models with chemotherapeutic drug-sensitive and -resistant clones to study the tumor clone dynamic. Our data demonstrated a competition between the drug-sensitive and drug-resistant clones, where the former exerted a suppressive effect on the proliferation of the latter in the absence of treatment. Our results also showed that metronomic schedule leads to better control of NSCLC cell heterogeneity than the standard-of-care MTD-like treatment schedule. Previous studies have demonstrated similar results in an ovarian and breast models (5,26). Indeed, their data showed that adaptive therapy strategies, which vary both the drug and dose density depending on the status of the tumor, maintained a stable population of drug-sensitive cells to suppress growth of resistant clones through intratumoral competition. Moreover, the emergence of targeted therapies, especially in NSCLC, has led to new schedule protocols with continuous drug administration as metronomic treatment do (27). Our study then supports the growing body of evidence that MTD-like treatment schedule might not be the optimal strategy to manage intratumor heterogeneity and points out the need to re-examine the current dogma of MTD.

With many effective therapies now available, it becomes more and more difficult to determine the optimal combinatorial treatments and schedules resulting in an unprecedented number of failed clinical trials over the past few years (28,29). Mathematical modeling has rapidly evolved and could provide relevant tools in oncology (30,31). Indeed, clinical trials in which mathematical models have be used to estimate the mechanism(s) of treatment failure and explore alternative strategies with new drugs and schedules showed the potential of mathematical modeling to improve patient outcomes (32–34). Here, our *in silico* prediction revealed that MC was the most suitable treatment schedule to manage NSCLC intratumor heterogeneity. This was furthermore confirmed in 3D spheroid and mouse xenografted model, supporting the interest of mathematical models to predict optimal treatments. Moreover, our models were also able to predict the clonal competition between drug-sensitive and -resistant clones. The clonal intratumor dynamic has led several researchers to develop modeling and simulation tools for describing biological and pharmacodynamics processes and aiming at managing resistance phenomena (5,35). More work must be done in the future to combine both types of models (*i.e* cancer clone dynamics with treatment schedule models) and design optimal treatment strategies to manage intratumor heterogeneity.

All tumors share a common phenotype of uncontrolled cell proliferation. In evolutionary cancer treatment, a key component of the Darwinian dynamics is the cost of resistance. Cancer cells must alter their phenotype to become resistant (36). To support the synthesis of biomass components and to generate energy required for cellular growth, cancer cells have to reshape the regulatory and functional properties of their metabolic networks (37). Cancer cells preferably use aerobic glycolysis to generate energy, which has been recognized as a hallmark of cancer (38). In our study, we demonstrated that drug-sensitive NSCLC clones mostly relied on aerobic glycolysis, while drug-resistant clones on OXPHOS activity. This observation is consistent with other studies reporting resistant cells being more addicted to mitochondrial functions in different cancer models including melanoma, leukemia and pancreatic cancers (39–41). Our results also revealed for the first time that metabolic activity of drug-sensitive clone could play a key role in controlling the proliferation of drug-resistant clones through a paracrine fashion. Whereas the role of extracellular vesicles as novel drug resistance modulators has emerged (16), our results showed an EV-independent mechanism involved in our NSCLC co-culture models. We highlighted that drug-sensitive cells exhibited higher glucose consumption and lactate production than drug-resistant cells. Lactate dehydrogenase A (LDHA) is a key glycolytic enzyme, a hallmark of aggressive cancers, and believed to be the major enzyme responsible for pyruvate to lactate conversion (19). Thus, inhibition of this key metabolic enzyme has been demonstrated as a promising strategy for cancer treatment (42,43). Our data however suggest that LDHA activity could play a major role in the regulation of drug-resistant clone proliferation. Inhibiting its activity might then lead to the rise of resistant clones. To support this hypothesis, more work is required to better understand the role of LDHA in the cancer cell clone heterogeneity.

By developing a 2D and 3D co-culture system of NSCLC, our results demonstrated that drug-sensitive clones exerted a suppressive effect on the growth of drug-resistant clones. We also revealed for the first time that this mechanism could involve a paracrine signaling that dependents on lactate dehydrogenase activity. Moreover, using mathematical modeling of clone expansion and response to treatment, our study highlighted metronomic schedule as the most effective strategy to achieve a long-term reduction of tumor volume as well as to better manage the intratumor heterogeneity. Altogether our study allows us to gain insights into the mechanism by which clones impact on each other and could open new therapeutic avenue to manage intratumor heterogeneity in NSCLC.

## Author contributions

**Conception and design:** EP, NA and MC

**Development of methodology:** MPM, GC, CC

**Acquisition of data:** MB, MLG, YS, ZR, MPM, MR, GC and CC

**Analysis and interpretation of data**: YS, ZR, MLG, GC and CC

**Writing the manuscript:** MB and MLG

**Study supervision:** MC and NA

**Other (funding acquisition):** EP, MC and NA.

All authors read the manuscript and made comments in order to improve it.

## Acknowledgments

This work was supported by grants from the not-for-profit organization ‘LN la Vie’, ‘Les Copains de Charles’, ‘RESOP’ and ‘Fondation Mont Ventoux’. This work has been carried out thanks to the support of ITMO Cancer. MB received support from AP-HM (Marseille, France). The authors wish to thank Laurence Borge, Victoire Gouirand and Sophie Vasseur for Seahorse and YSI analysis as well as Assia Benabdallah for mathematical modeling interpretations.

## Supplementary figures

**Figure S1.**
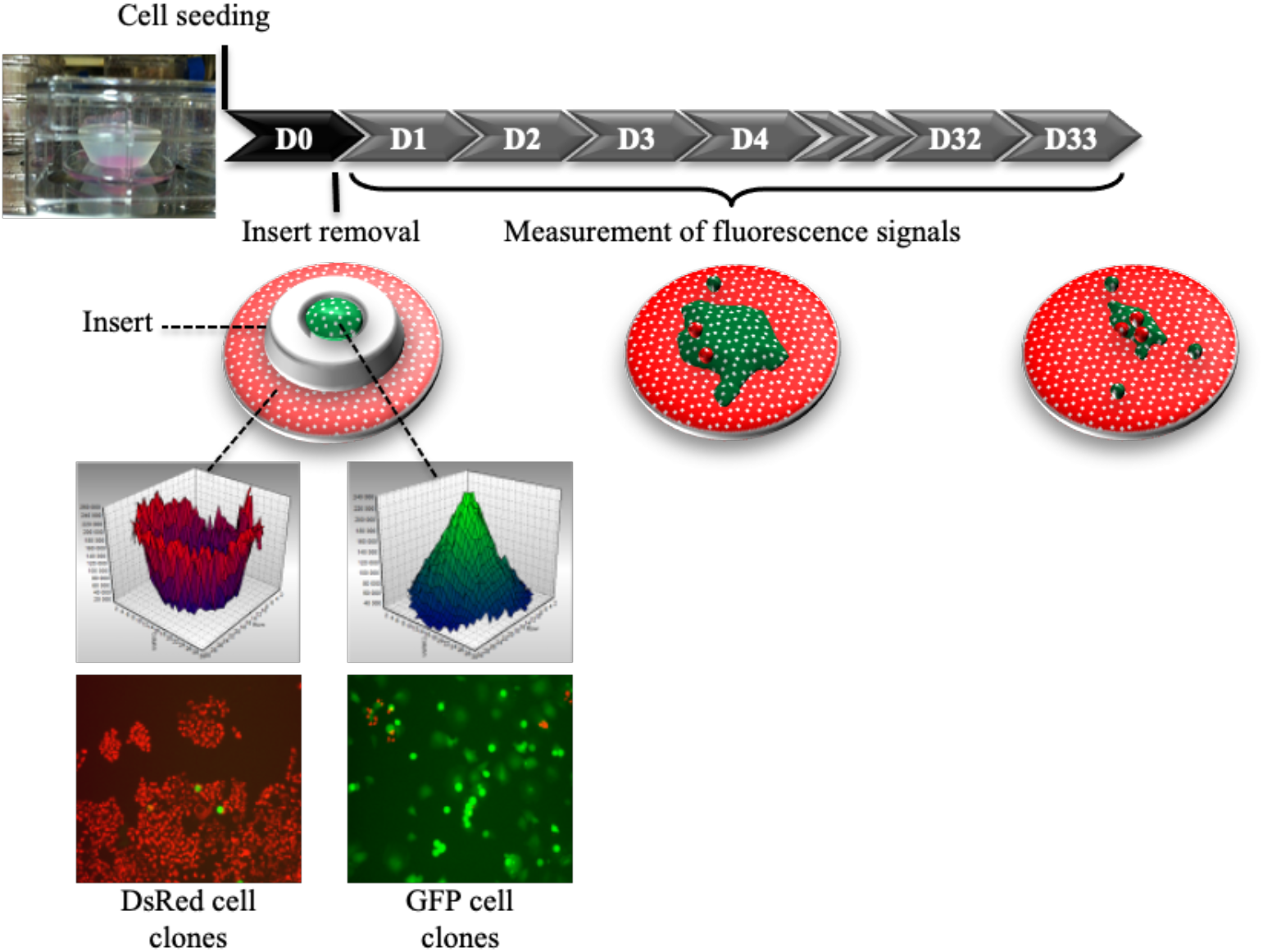
Schematic representation of heterogeneous adherent cell co-cultures using DsRed and GFP-expressing cells.

**Figure S2.**
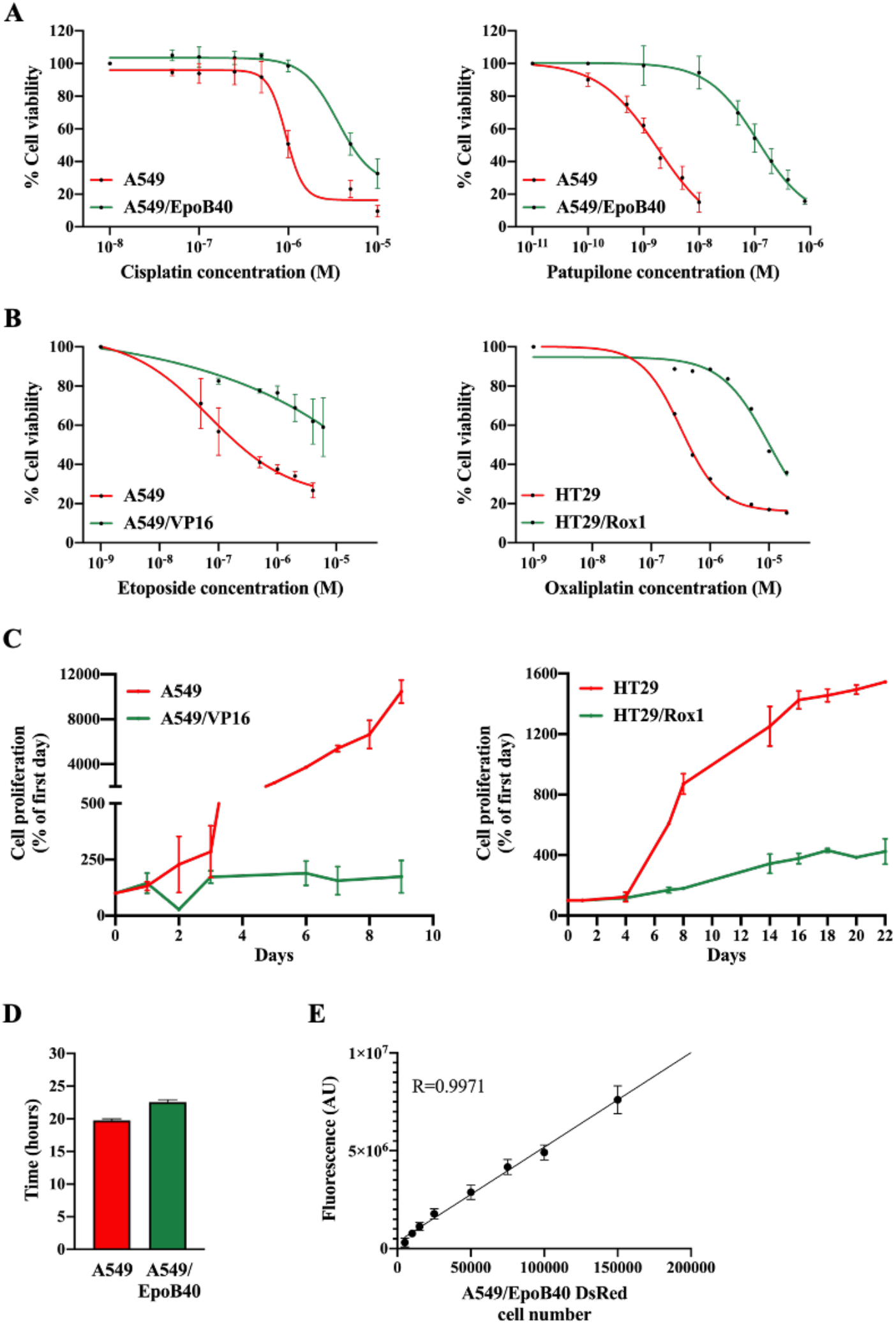
(A) Cell viability of A549 and A549/EpoB40 cells measured by MTT assay after a 72h exposition to increasing concentrations of chemotherapeutic agents (cisplatin or patupilone). Results were expressed as a percentage of viability in non-treated cells. (B) Cell viability of A549 and A549/VP16 cells (left panel) and HT29 and HT29/Roxl cells (right panel) measured by MTT assay after a 72h exposition to increasing concentrations of etoposide or oxaliplatin. Results were expressed as a percentage of viability in non-treated cells. (C) Cell proliferation of A549 and A549/VP16 (left panel) and HT29 and HT29/Roxl (right panel) by recording fluorescent signals (DsRed and GFP) over time. Data were expressed as a percentage of cell proliferation from day 0. (D) Doubling time of A549 and A549/EpoB40 cells assessed by real time impedance-based method. (E) Standard curve showing the conversion of fluorescent signal into a number of DsRed-expressing A549/EpoB40 cells in 3D co-culture models.

**Figure S3.**
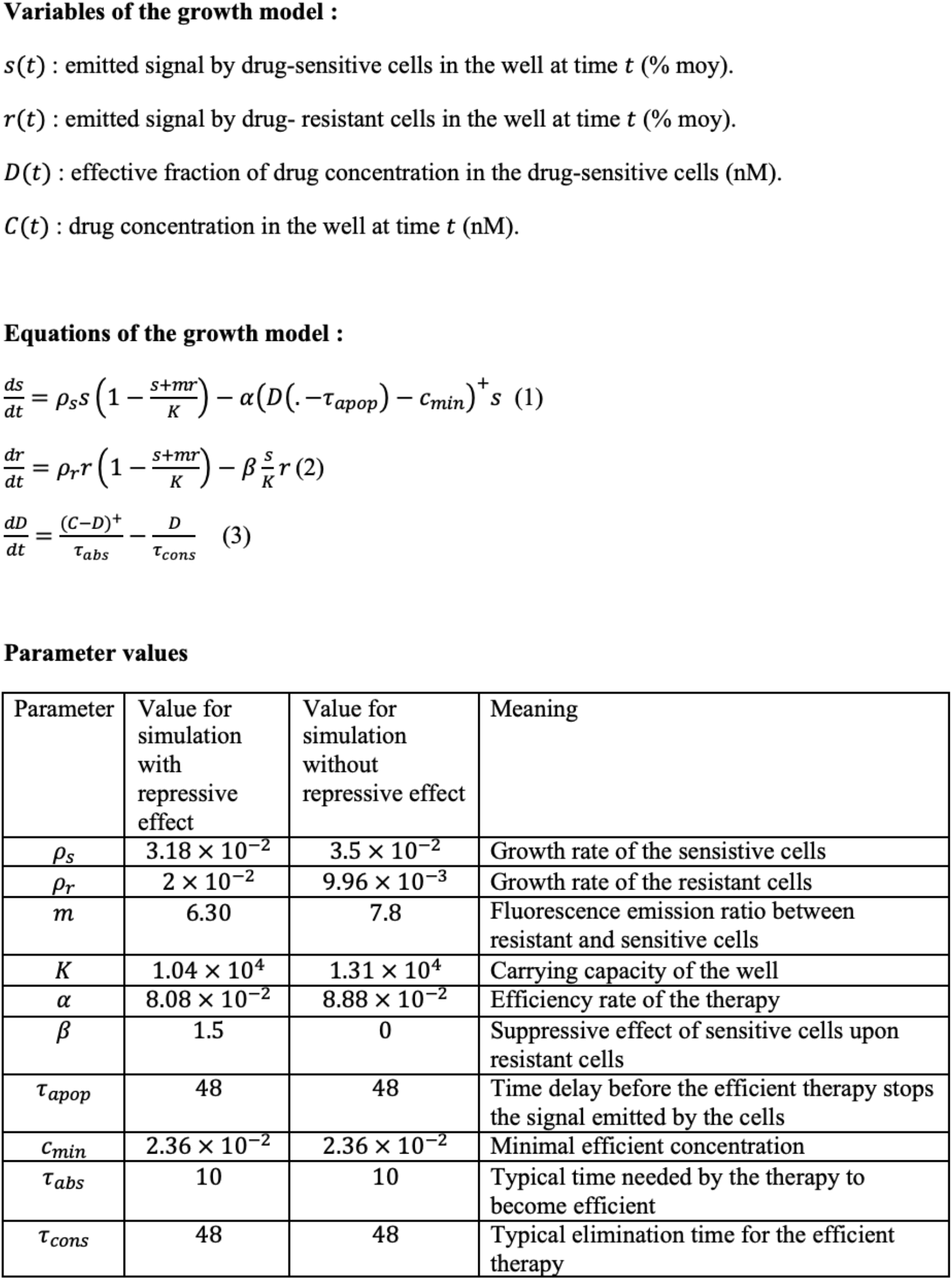
The mathematical model is described by a set of three equations where variables of the growth model and parameters values are detailed.

**Figure S4.**
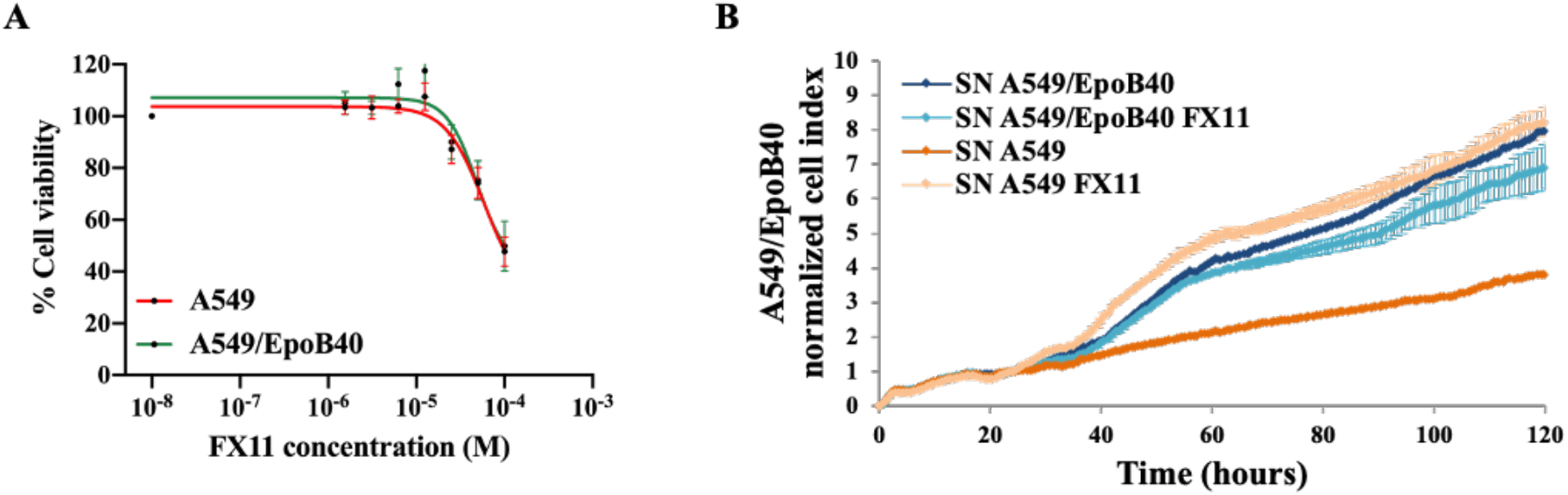
(A) Cell viability of A549 and A549/EpoB40 cells measured by MTT assay after a 72h exposition to increasing concentrations of FXII. Results were expressed as a percentage of viability in non-treated cells. (B) A549/EpoB40 cell growth was studied by real time impedance-based method. A549/EpoB40 cells were treated daily with cell-free supernatants from A549 cultures and from A549/EpoB40 cultures with or without 3μM of FXII. Measurements were performed every 15 minutes.

## References

1. Greaves M, Maley CC. Clonal evolution in cancer. Nature. 2012 Jan 18;481(7381):306–13.

2. Marusyk A, Almendro V, Polyak K. Intra-tumour heterogeneity: a looking glass for cancer? Nat Rev Cancer. 2012 Apr 19;12(5):323–34.

3. Dagogo-Jack I, Shaw AT. Tumour heterogeneity and resistance to cancer therapies. Nat Rev Clin Oncol. 2018;15(2):81–94.

4. Swanton C. Intratumor heterogeneity: evolution through space and time. Cancer Res. 2012 Oct 1;72(19):4875–82.

5. Gatenby RA. A change of strategy in the war on cancer. Nature. 2009 May 28;459(7246):508–9.

6. Gerlinger M, Swanton C. How Darwinian models inform therapeutic failure initiated by clonal heterogeneity in cancer medicine. Br J Cancer. 2010 Oct 12;103(8):1139–43.

7. Enriquez-Navas PM, Wojtkowiak JW, Gatenby RA. Application of Evolutionary Principles to Cancer Therapy. Cancer Res. 2015 Nov 15;75(22):4675–80.

8. Pasquier E, Kavallaris M, André N. Metronomic chemotherapy: new rationale for new directions. Nat Rev Clin Oncol. 2010 Aug;7(8):455–65.

9. André N, Carré M, Pasquier E. Metronomics: towards personalized chemotherapy? Nat Rev Clin Oncol. 2014 Jul;11(7):413–31.

10. Simsek C, Esin E, Yalcin S. Metronomic Chemotherapy: A Systematic Review of the Literature and Clinical Experience. J Oncol [Internet]. 2019 Mar 20 [cited 2020 Nov 28];2019. Available from: https://www.ncbi.nlm.nih.gov/pmc/articles/PMC6446118/

11. Savry A, Carre M, Berges R, Rovini A, Pobel I, Chacon C, et al. Bcl-2-enhanced efficacy of microtubule-targeting chemotherapy through Bim overexpression: implications for cancer treatment. Neoplasia N Y N. 2013 Jan;15(1):49–60.

12. Le Grand M, Berges R, Pasquier E, Montero M-P, Borge L, Carrier A, et al. Akt targeting as a strategy to boost chemotherapy efficacy in non-small cell lung cancer through metabolism suppression. Sci Rep. 2017 23;7:45136.

13. Timaner M, Beyar-Katz O, Shaked Y. Analysis of the Stromal Cellular Components of the Solid Tumor Microenvironment Using Flow Cytometry. Curr Protoc Cell Biol. 2016 Mar 1;70:19.18.1–19.18.12.

14. McGranahan N, Swanton C. Clonal Heterogeneity and Tumor Evolution: Past, Present, and the Future. Cell. 2017 09;168(4):613–28.

15. Calvo F, Sahai E. Cell communication networks in cancer invasion. Curr Opin Cell Biol. 2011 Oct;23(5):621–9.

16. Maacha S, Bhat AA, Jimenez L, Raza A, Haris M, Uddin S, et al. Extracellular vesicles-mediated intercellular communication: roles in the tumor microenvironment and anti-cancer drug resistance. Mol Cancer. 2019 30;18(1):55.

17. Brinton LT, Sloane HS, Kester M, Kelly KA. Formation and role of exosomes in cancer. Cell Mol Life Sci CMLS. 2015 Feb;72(4):659–71.

18. Yoshida GJ. Metabolic reprogramming: the emerging concept and associated therapeutic strategies. J Exp Clin Cancer Res CR. 2015 Oct 6;34:111.

19. Mishra D, Banerjee D. Lactate Dehydrogenases as Metabolic Links between Tumor and Stroma in the Tumor Microenvironment. Cancers. 2019 May 29;11(6).

20. Cui J, Shi M, Xie D, Wei D, Jia Z, Zheng S, et al. FOXM1 promotes the warburg effect and pancreatic cancer progression via transactivation of LDHA expression. Clin Cancer Res Off J Am Assoc Cancer Res. 2014 May 15;20(10):2595–606.

21. Jiang F, Ma S, Xue Y, Hou J, Zhang Y. LDH-A promotes malignant progression via activation of epithelial-to-mesenchymal transition and conferring stemness in muscle-invasive bladder cancer. Biochem Biophys Res Commun. 2016 Jan 22;469(4):985–92.

22. Torre LA, Bray F, Siegel RL, Ferlay J, Lortet-Tieulent J, Jemal A. Global cancer statistics, 2012. CA Cancer J Clin. 2015 Mar;65(2):87–108.

23. Herbst RS, Morgensztern D, Boshoff C. The biology and management of non-small cell lung cancer. Nature. 2018 24;553(7689):446–54.

24. Jamal-Hanjani M, Wilson GA, McGranahan N, Birkbak NJ, Watkins TBK Veeriah S, et al. Tracking the Evolution of Non-Small-Cell Lung Cancer. N Engl J Med. 2017 01;376(22):2109–21.

25. Goto T, Hirotsu Y, Amemiya K, Mochizuki H, Omata M. Understanding Intratumor Heterogeneity and Evolution in NSCLC and Potential New Therapeutic Approach. Cancers. 2018 Jun 22;10(7).

26. Enriquez-Navas PM, Kam Y, Das T, Hassan S, Silva A, Foroutan P, et al. Exploiting evolutionary principles to prolong tumor control in preclinical models of breast cancer. Sci Transl Med. 2016 Feb 24;8(327):327ra24.

27. Hirsch FR, Scagliotti GV, Mulshine JL, Kwon R, Curran WJ, Wu Y-L, et al. Lung cancer: current therapies and new targeted treatments. Lancet Lond Engl. 2017 21;389(10066):299–311.

28. Prasad V, De Jesús K, Mailankody S. The high price of anticancer drugs: origins, implications, barriers, solutions. Nat Rev Clin Oncol. 2017 Jun;14(6):381–90.

29. Jardim DL, de Melo Gagliato D, Giles FJ, Kurzrock R. Analysis of Drug Development Paradigms for Immune Checkpoint Inhibitors. Clin Cancer Res Off J Am Assoc Cancer Res. 2018 15;24(8):1785–94.

30. Rockne RC, Hawkins-Daarud A, Swanson KR, Sluka JP, Glazier JA, Macklin P, et al. The 2019 mathematical oncology roadmap. Phys Biol. 2019 19;16(4):041005.

31. Benzekry S. Artificial Intelligence and Mechanistic Modeling for Clinical Decision Making in Oncology. Clin Pharmacol Ther. 2020 Jun 18;

32. Barlesi F, Imbs D-C, Tomasini P, Greillier L, Galloux M, Testot-Ferry A, et al. Mathematical modeling for Phase I cancer trials: A study of metronomic vinorelbine for advanced non-small cell lung cancer (NSCLC) and mesothelioma patients. Oncotarget. 2017 Jul 18;8(29):47161–6.

33. West JB, Dinh MN, Brown JS, Zhang J, Anderson AR, Gatenby RA. Multidrug Cancer Therapy in Metastatic Castrate-Resistant Prostate Cancer: An Evolution-Based Strategy. Clin Cancer Res Off J Am Assoc Cancer Res. 2019 15;25(14):4413–21.

34. Zhang J, Cunningham JJ, Brown JS, Gatenby RA. Integrating evolutionary dynamics into treatment of metastatic castrate-resistant prostate cancer. Nat Commun. 2017 28;8(1):1816.

35. Chmielecki J, Foo J, Oxnard GR, Hutchinson K, Ohashi K, Somwar R, et al. Optimization of dosing for EGFR-mutant non-small cell lung cancer with evolutionary cancer modeling. Sci Transl Med. 2011 Jul 6;3(90):90ra59.

36. Stanková K, Brown JS, Dalton WS, Gatenby RA. Optimizing Cancer Treatment Using Game Theory: A Review. JAMA Oncol. 2019 01;5(1):96–103.

37. Guerra F, Arbini AA, Moro L. Mitochondria and cancer chemoresistance. Biochim Biophys Acta Bioenerg. 2017 Aug;1858(8):686–99.

38. Hanahan D, Weinberg RA. Hallmarks of cancer: the next generation. Cell. 2011 Mar 4;144(5):646–74.

39. Haq R, Shoag J, Andreu-Perez P, Yokoyama S, Edelman H, Rowe GC, et al. Oncogenic BRAF regulates oxidative metabolism via PGC1α and MITF. Cancer Cell. 2013 Mar 18;23(3):302–15.

40. Viale A, Pettazzoni P, Lyssiotis CA, Ying H, Sánchez N, Marchesini M, et al. Oncogene ablation-resistant pancreatic cancer cells depend on mitochondrial function. Nature. 2014 Oct 30;514(7524):628–32.

41. Farge T, Saland E, de Toni F, Aroua N, Hosseini M, Perry R, et al. Chemotherapy-Resistant Human Acute Myeloid Leukemia Cells Are Not Enriched for Leukemic Stem Cells but Require Oxidative Metabolism. Cancer Discov. 2017;7(7):716–35.

42. Fantin VR, St-Pierre J, Leder P. Attenuation of LDH-A expression uncovers a link between glycolysis, mitochondrial physiology, and tumor maintenance. Cancer Cell. 2006 Jun;9(6):425–34.

43. Wang Z-Y, Loo TY, Shen J-G, Wang N, Wang D-M, Yang D-P, et al. LDH-A silencing suppresses breast cancer tumorigenicity through induction of oxidative stress mediated mitochondrial pathway apoptosis. Breast Cancer Res Treat. 2012 Feb;131(3):791–800.

